# Identification and distribution of novel candidate T6SS effectors encoded in *Salmonella* Pathogenicity island 6

**DOI:** 10.1101/2023.07.03.547122

**Authors:** Carlos J. Blondel, Fernando A. Amaya, Paloma Bustamante, Carlos A. Santiviago, David Pezoa

## Abstract

The type VI secretion system (T6SS) is a contact-dependent contractile multiprotein apparatus widely distributed in Gram-negative bacteria. These systems can deliver different effector proteins into target bacterial and/or eukaryotic cells, contributing to the environmental fitness and virulence of many bacterial pathogens. *Salmonella* harbors five different T6SSs encoded in different genomic islands. The T6SS encoded in *Salmonella* Pathogenicity Island 6 (SPI-6) contributes to *Salmonella* competition with the host microbiota and its interaction with infected host cells. Despite its relevance, information regarding the total number of effector proteins encoded within SPI-6 and its distribution among different *Salmonella enterica* serotypes is limited. In this work, we performed bioinformatic and comparative genomics analyses of the SPI-6 T6SS gene cluster to expand our knowledge regarding the T6SS effector repertoire and the global distribution of these effectors in *Salmonella*. The analysis of a curated dataset of 60 *Salmonella enterica* genomes from the Secret6 database revealed the presence of 23 novel putative T6SS effector/immunity protein (E/I) modules. These effectors were concentrated in the variable regions 1 to 3 (VR1-3) of the SPI-6 T6SS gene cluster. VR1-2 were enriched in candidate effectors with predicted peptidoglycan hydrolase activity, while VR3 was enriched in candidate effectors of the Rhs family with C-terminal extensions with predicted DNase, RNase, deaminase, or ADP-ribosyltransferase activity. A global analysis of known and candidate effector proteins in *Salmonella enterica* genomes from the NCBI database revealed that T6SS effector proteins are differentially distributed among *Salmonella* serotypes. While some effectors are present in over 200 serotypes, others are found in less than a dozen. A hierarchical clustering analysis identified *Salmonella* serotypes with distinct profiles of T6SS effectors and candidate effectors, highlighting the diversity of T6SS effector repertoires in *Salmonella enterica*. The existence of different repertoires of effector proteins suggests that different effector protein combinations may have a differential impact on the environmental fitness and pathogenic potential of these strains.

## INTRODUCTION

The type VI secretion system (T6SS) is a multiprotein nanomachine composed of 13 structural components and various accessory proteins that deliver protein effectors into target cells through a contractile mechanism (Zoued *et al*., 2014; Ho *et al*., 2014; Coulthurst, 2019). The T6SS needle, composed of an inner tube (made of Hcp hexamers) tipped by a spike complex comprising the VgrG and PAAR proteins, is wrapped into a contractile sheath formed by the polymerization of TssB/TssC subunits. These are assembled into an extended, metastable conformation (Silverman *et al*., 2013). Contraction of the sheath upon contact with a target cell or sensing cell envelope damage propels the needle toward the target cell (Brackmann *et al*., 2017). T6SS effector proteins are classified as either cargo or specialized effectors. Cargo effectors are delivered by either non-covalent interaction with some core components (Coulthurst, 2019), while specialized effectors are often fused to either VgrG, Hcp or PAAR proteins (Durand *et al*., 2014; Whitney *et al*., 2014; Diniz and Coulthurst, 2015; Ma *et al*., 2017; Pissaridou *et al*., 2018).

The extensive repertoire of effector proteins makes the T6SS a highly versatile machine that can target prokaryotic or eukaryotic cells (Ma and Mekalanos, 2010; Russell *et al*., 2012, 2013; Koskiniemi *et al*., 2014; Miyata *et al*., 2013; Srikannathasan *et al*., 2013; Whitney *et al*., 2013; Egan *et al*., 2015; Bondage *et al*., 2016; Flaugnatti *et al*., 2016; Tang *et al*., 2018; Ting *et al*., 2018; Ahmad *et al*., 2019; Berni *et al*., 2019; Coulthurst, 2019; Jana *et al*., 2019; Mariano *et al*., 2019; Wood *et al*., 2020). Among the anti-bacterial effector proteins, some target the peptidic or glycosidic bonds of the peptidoglycan (Ma and Mekalanos, 2010; Russell *et al*., 2012; Srikannathasan *et al*., 2013; Whitney *et al*., 2013; Berni *et al*., 2019; Wood *et al*., 2019), or the FtsZ cell division ring (Ting *et al*., 2018). These anti-bacterial effectors are encoded in bi-cistronic elements with their cognate immunity proteins (effector/immunity pairs) that bind tightly and specifically to their cognate effector preventing self-intoxication and killing of sibling cells (Russell *et al*., 2012). Other T6SS effectors are eukaryote-specific, such as those targeting the actin or microtubule cytoskeleton networks (Pukatzki *et al*., 2007; Ma *et al*., 2009; Ma and Mekalanos, 2010; Miyata *et al*., 2011; Zheng *et al*., 2011; Durand *et al*., 2012; Lindgren *et al*., 2013; Schwarz *et al*., 2014; Heisler *et al*., 2015; Sana *et al*., 2015; Aubert *et al*., 2016; Jiang *et al*., 2016; Ray *et al*., 2017; Dutta *et al*., 2019; Tan *et al*., 2019; Wood *et al*., 2019), and others can target both bacterial and eukaryotic cells (trans-kingdom effectors). These effectors include those targeting conserved molecules (NAD) and macromolecules (DNA, phospholipids) or forming pores in membranes (Whitney *et al*., 2015; Tang *et al*., 2018; Ahmad *et al*., 2019).

Many enteric pathogens (e.g., *Salmonella*, *Shigella* and *Vibrio*) use the T6SS to colonize the intestinal tract of infected hosts (Sana *et al*., 2016; Chassaing *et al*., 2018), while some of the commensal *Bacteroides* strains of the gut use their T6SSs for protection against pathogens (Coyne and Comstock, 2019). The T6SS is, therefore, a key player in bacterial warfare.

Salmonellosis is a foodborne bacterial disease caused by different serotypes of *Salmonella enterica* (GBD-2017 *et al*., 2019). Worldwide, this illness is responsible for 95.1 million cases of gastroenteritis per year (GBD-2017 *et al*., 2019). *Salmonella* genus includes more than 2,600 serotypes distributed between species *S*. *enterica* and *S*. *bongori* (Issenhuth-Jeanjean *et al*., 2014), which differ in clinical signs and host range (Uzzau *et al*., 2000). In addition, the World Health Organization (WHO) has also included *Salmonella* as a high-priority pathogen due to the emergence of strains with high levels of fluoroquinolone resistance (GBD-2017 *et al*., 2019). In *Salmonella*, 5 T6SS gene clusters have been identified within *Salmonella* Pathogenicity Islands (SPIs) SPI-6, SPI-19, SPI-20, SPI-21, and SPI-22 (Blondel *et al*., 2009; Fookes *et al*., 2011). These T6SSs are distributed in 4 different evolutionary lineages: T6SS_SPI-6_ belongs to subtype i3, T6SS_SPI-19_ to subtype i1, T6SS_SPI-22_ to subtype i4a, and both T6SS_SPI-20_ and T6SS_SPI-21_ belong to subtype i2 (Bao *et al*., 2019). Besides their distinct evolutionary origin, these five T6SS gene clusters are differentially distributed among distinct serotypes, subspecies, and species of *Salmonella* (Blondel *et al*., 2009).

Notably, most of these T6SSs have been shown to contribute to the virulence and pathogenesis of different *Salmonella* serotypes (Mulder *et al*., 2012, Pezoa *et al*., 2013, Pezoa *et al*., 2014, Sana *et al*., 2016, Sibinelli-Sousa *et al*., 2022, Xian *et al*., 2020, Blondel *et al*., 2010; Hespanhol *et al*., 2022). One of the most studied and widely distributed T6SS corresponds to that encoded in SPI-6. Depending on the serotype, the SPI-6 T6SS gene cluster comprises a region of ∼35 to 50 kb encoding ∼30 to 45 ORFs, including each of the 13 T6SS core components. The genetic architecture of the SPI-6 T6SS gene cluster is highly conserved among serotypes; nonetheless, there are structural differences restricted to three variable regions of the island (herein referred to as VR1, VR2, and VR3, **Figure 1**) (Blondel *et al*., 2009). In *S.* Typhimurium and *S*. Dublin, 9 SPI-6 T6SS effector proteins have been described to date (Russell *et al*., 2012; Benz *et al*., 2013; Sana *et al*., 2016; Whitney *et al*., 2013; Sibinelli-Sousa *et al*., 2020; Lorente-Cobo *et al*., 2022; Koskiniemi *et al*., 2014; Amaya *et al*., 2022; Jurenas *et al*., 2022), most of which are encoded within these variable regions (**Figure 1**).

**Figure 1.**
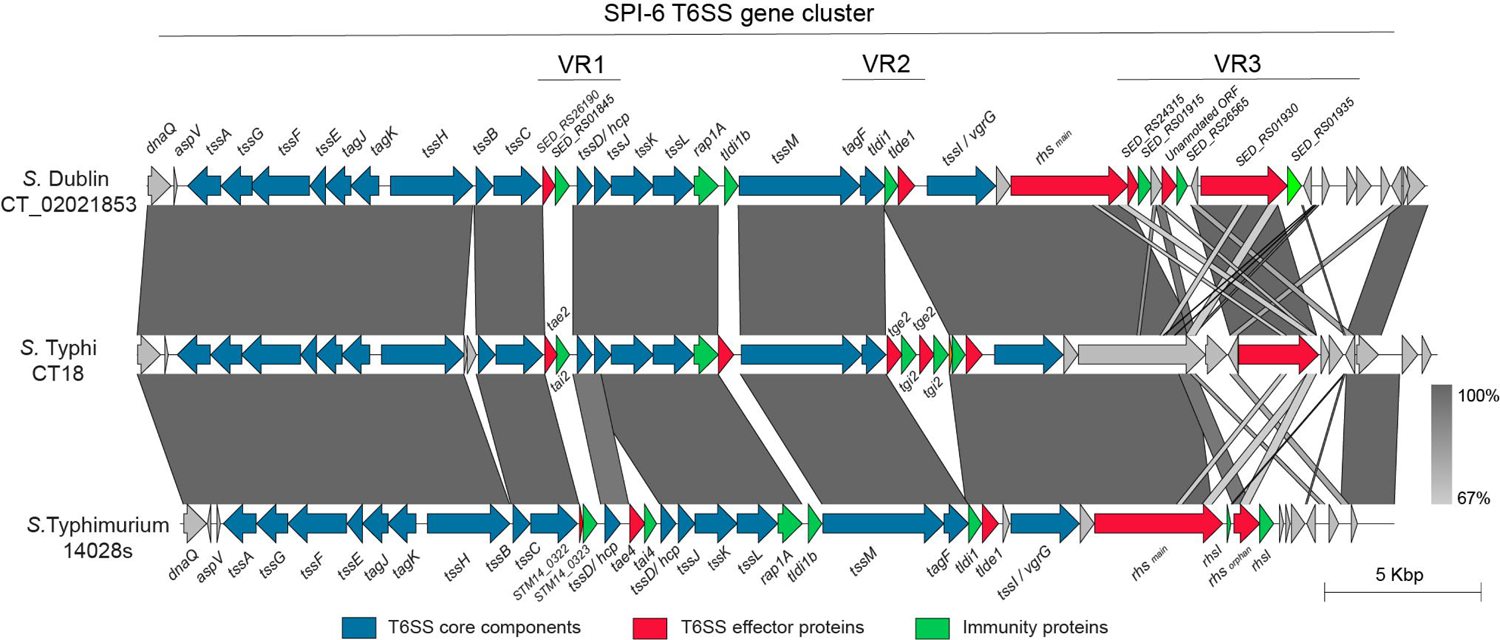
Schematic representation of selected SPI-6 T6SS gene clusters. The figure shows an alignment of the T6SS_SPI-6_ gene cluster of *S*. Dublin CT_02021853, *S*. Typhi CT18 and *S*. Typhimurium 14028s. The location of variable regions 1 to 3 is shown. ORF encoding previously described T6SS effectors and cognate immunity proteins are shown in red and green, respectively. ORF encoding T6SS core components are shown in blue. Grayscale represents the percentage of identity between nucleotide sequences.

The VR1 is located downstream of gene *tssC* and encodes the E/I modules Tae2/Tai2 and Tae4/Tai4. Tae2 and Tae4 are peptidoglycan hydrolases able to cleave the DD-crosslinks between D-mDAP and D-alanine or the covalent link between D-Glu and mDAP of the tetrapeptide stem, respectively, thus contributing to interbacterial competition and mice colonization (Russell *et al*., 2012; Sana *et al*., 2016). VR2 is located downstream of gene *tssM* and encodes many proteins of unknown function and two E/I modules with peptidoglycan hydrolase activity: Tge2/Tgi2P is predicted to have N-acetylglucosaminidase activity (Whitney *et al*., 2013), while Tlde1/Tldi shows L,D carboxypeptidase activity against the peptide stems of the peptidoglycan layer (Sibinelli-Sousa *et al*., 2020; Lorente-Cobo *et al*., 2022). Finally, the VR3 is located downstream of gene *tssI* and encodes a variable number of Rhs elements, some of them harboring endonuclease domains such as HNHc (DNase), Ntox47 (RNase) and an ART domain (ADP-ribosyltransferase) linked to the C-terminal of these Rhs proteins (Koskiniemi *et al*., 2014; Amaya *et al*., 2022; Jurenas *et al*., 2022).

Most of our knowledge regarding the presence and distribution of SPI-6 T6SS effector proteins comes from studies using reference strains of a limited number of serotypes (e.g., *S.* Typhimurium and *S.* Dublin) (Russell *et al*., 2012; Benz *et al*., 2013; Sana *et al*., 2016; Whitney *et al*., 2013; Sibinelli-Sousa *et al*., 2020; Lorente-Cobo *et al*., 2022; Koskiniemi *et al*., 2014; Amaya *et al*., 2022; Jurenas *et al*., 2022). In this study, we performed a bioinformatic prediction analysis searching for putative T6SS effectors in a dataset of 60 genomes covering 37 *S. enterica* serotypes retrieved from the curated Secret6 database. Our analysis identified 23 novel putative anti-bacterial effectors encoded in E/I modules within the 3 VRs of the SPI-6 T6SS gene cluster. These candidates include 5 effectors with putative peptidoglycan hydrolase activity, 16 effectors with potential nuclease activity and 2 effectors targeting bacterial protein translation. Finally, we expanded our analysis to include all available *Salmonella* genomes deposited in the NCBI database and determined the global distribution of these novel putative effectors. A hierarchical clustering analysis identified that some effectors are conserved in most *Salmonella* serotypes. In contrast, most other effectors are differentially distributed in different serotypes. The presence of different sets of T6SS effectors suggests that distinct repertoires of these proteins may have a differential impact on the pathogenicity and environmental adaptation of *Salmonella* serotypes.

## MATERIALS AND METHODS

### Identification of candidate SPI-6 T6SS effectors

First, we searched the Secret6 database (https://bioinfo-mml.sjtu.edu.cn/SecReT6/download.html) for *Salmonella* genomes encoding the minimal 13 core components of a T6SS and identified a total of 60 genomes that met this requirement. Then, to identify putative T6SS effectors encoded within SPI-6 of *Salmonella*, each ORF of this island was analyzed with the Bastion6 pipeline (Wang *et al*., 2018) excluding the 13 T6SS core components. ORFs presenting a Bastion6 score ≥ 0.7 were considered as candidate T6SS effectors. Each Bastion6 prediction was further analyzed with tools implemented in the Operon-Mapper web server (Taboada *et al*., 2018) to determine if it was part of a bi-cistronic unit also encoding a putative immunity protein [i.e., a small protein with potential signal peptides (SignalP 6.0) and/or transmembrane domains (TMHMM 2.0)]. Identification of conserved functional domains and motifs in the candidate T6SS effectors was performed using the PROSITE, NCBI-CDD, Motif-finder, and Pfam databases (Kanehisa *et al*., 2002; Sigrist *et al*., 2013; Finn *et al*., 2014; Lu *et al*., 2019) implemented in the GenomeNet search engine. An e-value cutoff score of 0.01 was used. Finally, a biochemical functional prediction for each putative effector and immunity protein identified was performed by HMM homology searches using the HHpred HMM-HMM comparison tool (Zimmermann *et al*., 2017).

### Hierarchical clustering analysis of the novel SPI-6 T6SS effectors

For hierarchical clustering analysis, a presence/absence matrix of each T6SS effector and candidate effector was constructed for each bacterial genome and uploaded as a csv file to the online server MORPHEUS (https://software.broadinstitute.org/morpheus) using default parameters (i.e., one minus Pearson’s correlation, average linkage method).

### *Salmonella* 16S rDNA phylogenetic analyses

The 16S rDNA sequences were obtained from the 60 *Salmonella* genomes previously analyzed. The sequences were concatenated and aligned with ClustalW using the Molecular Evolutionary Genetics Analysis (MEGA) software version 7.0 (Kumar *et al*., 2016). A phylogenetic tree was built from the alignments obtained from MEGA by performing a bootstrap test of phylogeny (1,000 replications) using the maximum-likelihood method with a Jones-Taylor-Thornton correction model.

### Sequence and phylogenetic analyses

Identification of the 23 novel T6SS effector orthologs was carried out using the DNA sequence of each E/I module in BLASTn analyses of all publicly available *Salmonella* genome sequences deposited in the NCBI database (October 2022). A 90% identity and 90% sequence coverage threshold was used to select positive matches. Sequence conservation was analyzed by multiple sequence alignments using MAFFT (Katoh *et al*., 2017) and T-Coffee Expresso (Notredame *et al*., 2000) and visualized by ESPript 3 (Robert and Gouet, 2014). Comparative genomic analysis of SPI-6 T6SS gene clusters was performed using the multiple genome alignment tool Mauve (Darling *et al*., 2004) and EasyFig v2.2.5 (Sullivan *et al*., 2011). Nucleotide sequences were analyzed by the sequence visualization and annotation tool Artemis version 18 (Rutherford *et al*., 2000).

## RESULTS

### Analysis of a curated dataset of *Salmonella* genomes reveals 23 novel putative E/I modules encoded within the SPI-6 T6SS gene cluster

To identify high-confidence novel T6SS effector, we first screened the SPI-6 T6SS gene clusters of a dataset of 60 *Salmonella enterica* genomes from the Secret6 curated database (Zhang *et al*., 2023). This database includes 60 strains covering 37 *Salmonella* serotypes (**Table S1**). Each ORF within SPI-6 T6SS gene clusters was analyzed based on four criteria: i) identification of candidate effectors through Bastion6 analysis (a bioinformatic tool that predicts T6SS effectors based on amino acid sequence, evolutionary information, and physicochemical properties); ii) identification of putative immunity proteins by detection of signal peptides (SignalP 6.0), transmembrane domains (TMHMM 2.0) and operon prediction (Operon-mapper; Taboada *et al*., 2018); iii) identification of conserved functional domains associated with *bona fide* T6SS effectors (INTERPROSCAN, PROSITE, NCBI-CDD, MOTIF, and Pfam) and iv) functional biochemical prediction using the HHpred HMM-HMM server. In addition, we analyzed these gene clusters to identify potential unannotated ORFs which could encode putative effectors and cognate immunity proteins.

Our analysis identified 23 novel putative effector proteins and cognate immunity proteins (**Table 1**). These candidates included both cargo and specialized effector proteins with diverse predicted biochemical functions, including peptidoglycan hydrolases (5), DNases (8), RNases (6), deaminases (1), ADP-ribosyltransferases (2) and hybrid DNases/RNases (1) (**Table 1**). In addition, our analysis showed that the repertoire of E/I modules in SPI-6 vary considerably between closely related strains (**Figure 2**). Of note, comparative genomic analyses revealed that each identified E/I module is encoded within one of the 3 VRs previously described (Blondel *et al*., 2009). One E/I module is encoded within VR1, four within VR2, and 18 are encoded within VR3 (**Figure 3**).

**Figure 2.**
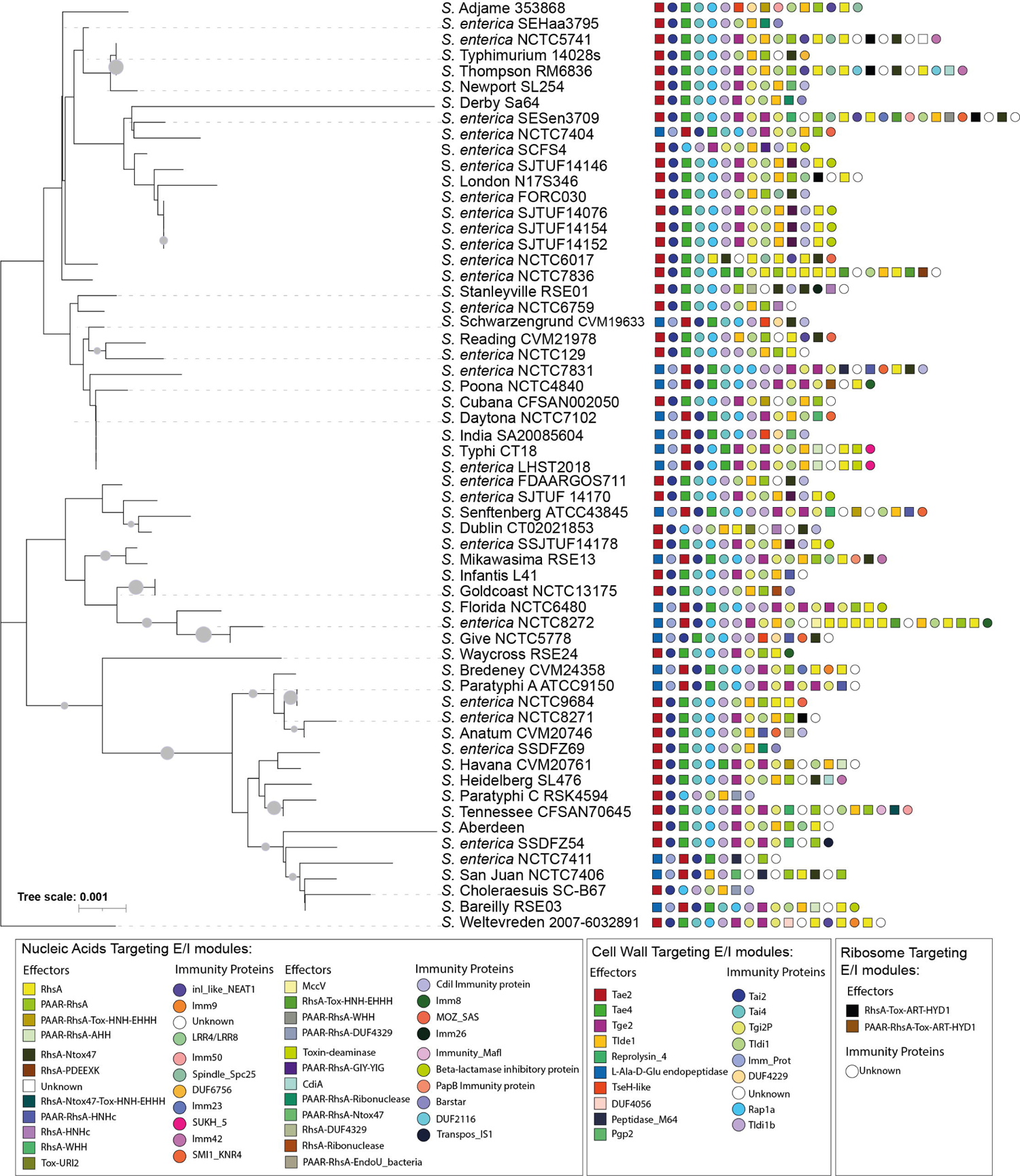
16S rDNA phylogeny and T6SS E/I module composition of *Salmonella enterica* SPI-6. Concatenated 16S rDNA nucleotide sequences from 60 *Salmonella* genomes deposited in Secret6 database were aligned with ClustalW using MEGA version 7.0. Next, a maximum-likelihood phylogenetic tree was built from the alignment using a bootstrap test of phylogeny (1,000 replications) with a Jones-Taylor-Thornton correction model. Squares and circles next to each strain name correspond to an effector or an immunity protein, respectively. Different colors represent confirmed or predicted functions, as indicated in the figure.

**Figure 3.**
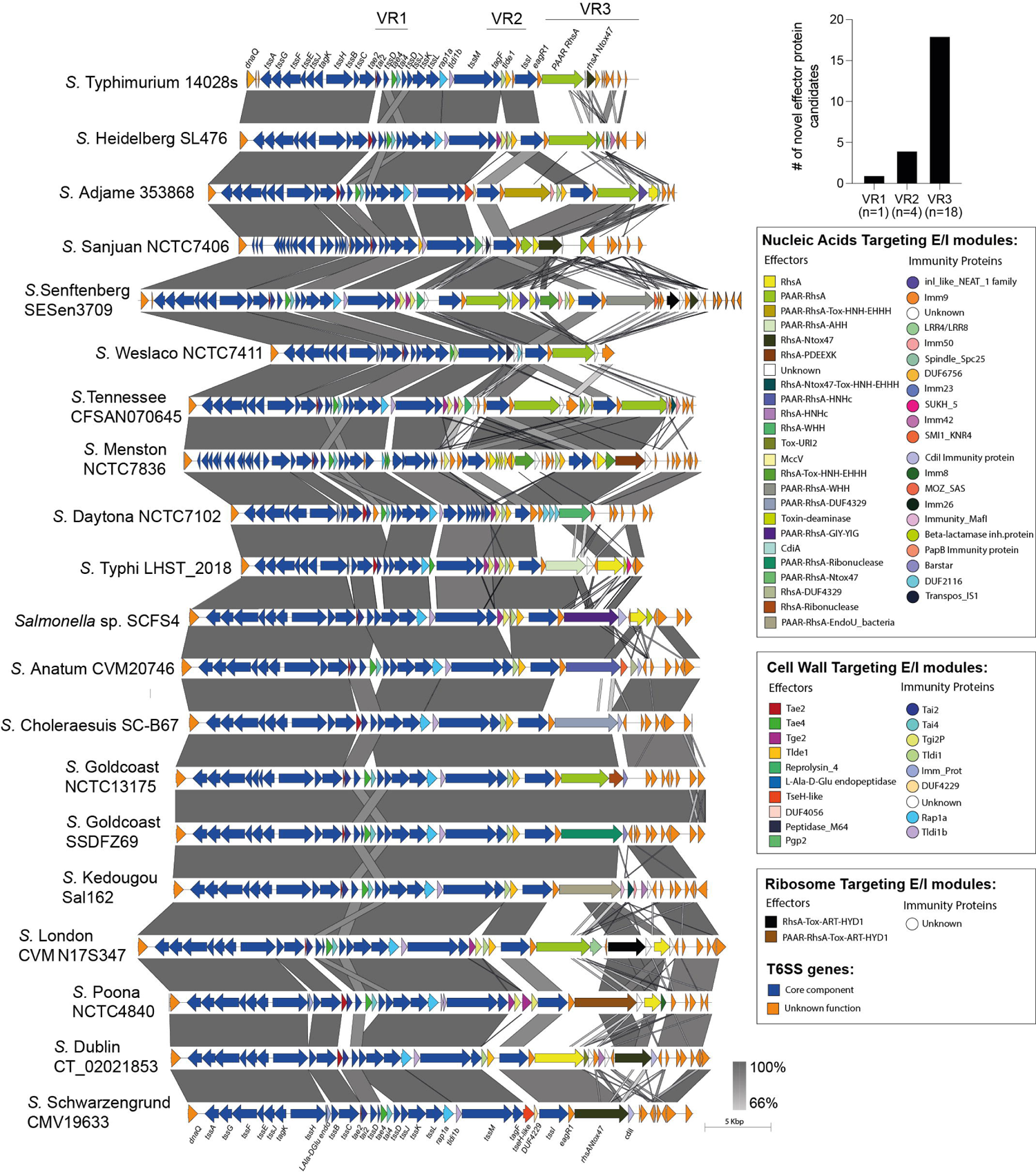
Comparative genomic analysis of SPI-6 T6SS gene clusters in representative *Salmonella* serotypes. The location of variable regions 1 to 3 is shown. ORFs encoding T6SS core components are shown in blue. ORFs encoding E/I modules are presented in different colors according to the confirmed or predicted functions, as indicated in the figure. The three variable regions of the T6SS_SPI-6_ gene cluster are demarked by black lines. Grayscale represents the percentage of identity between nucleotide sequences.

**Table 1.**
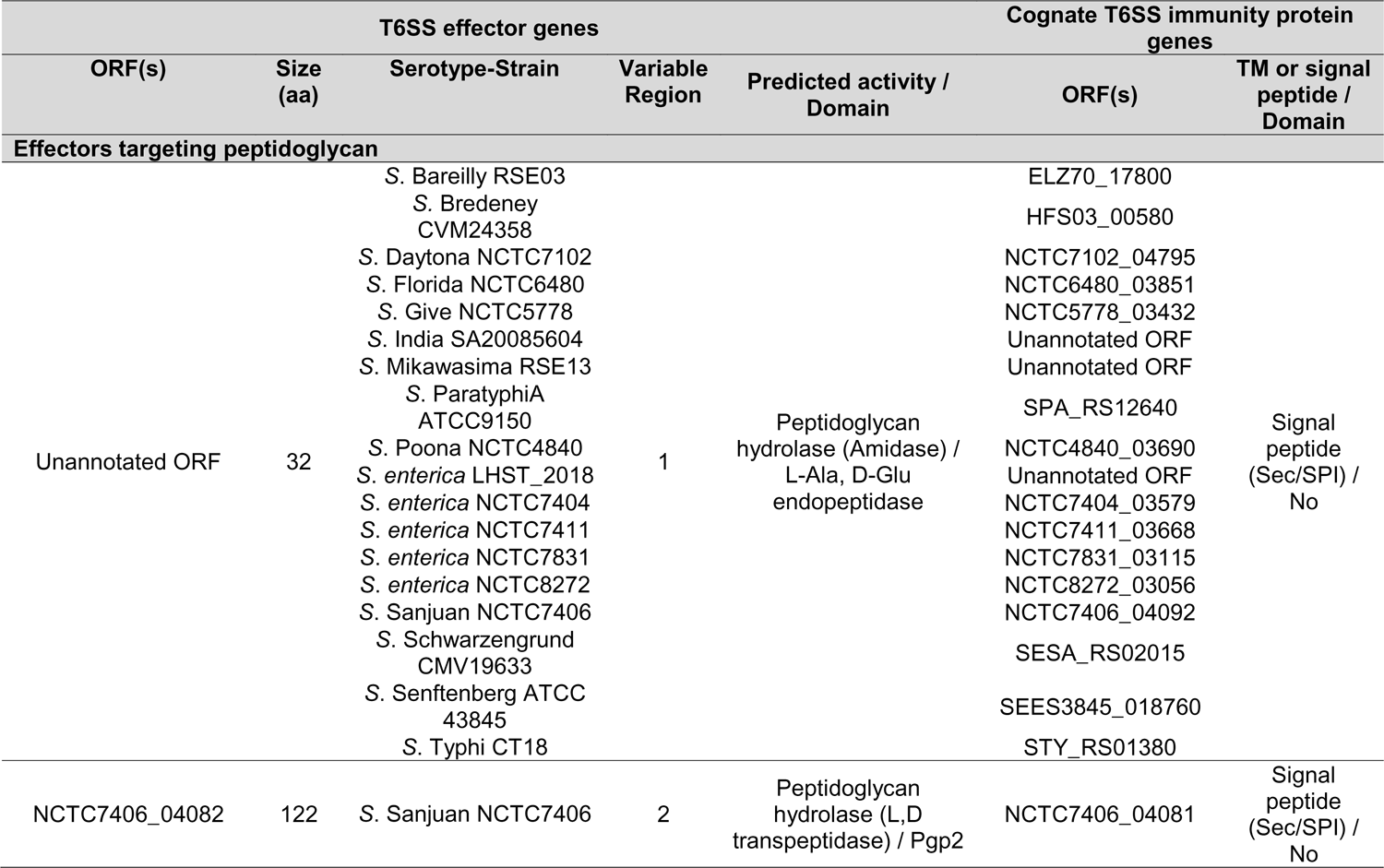
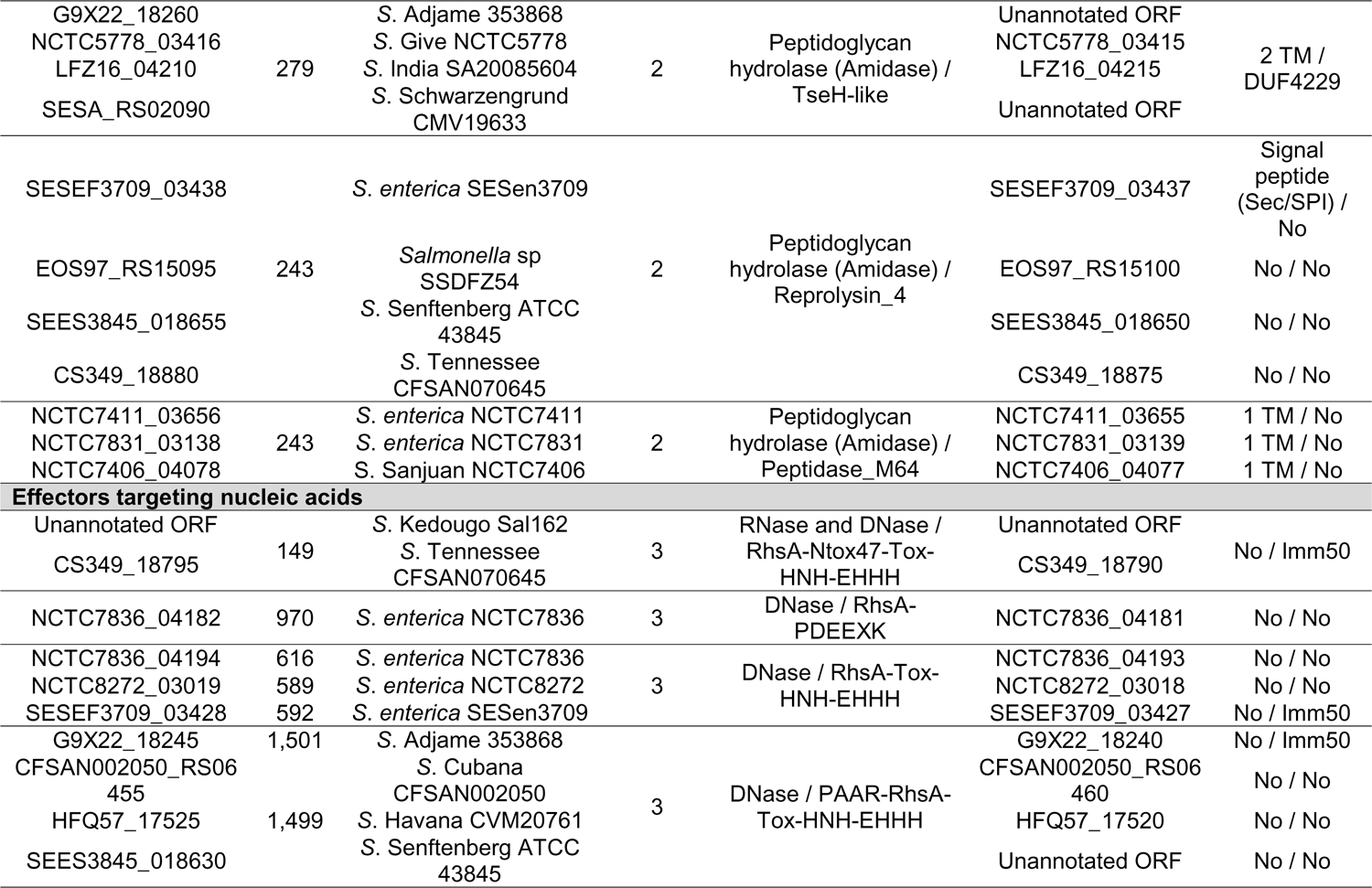
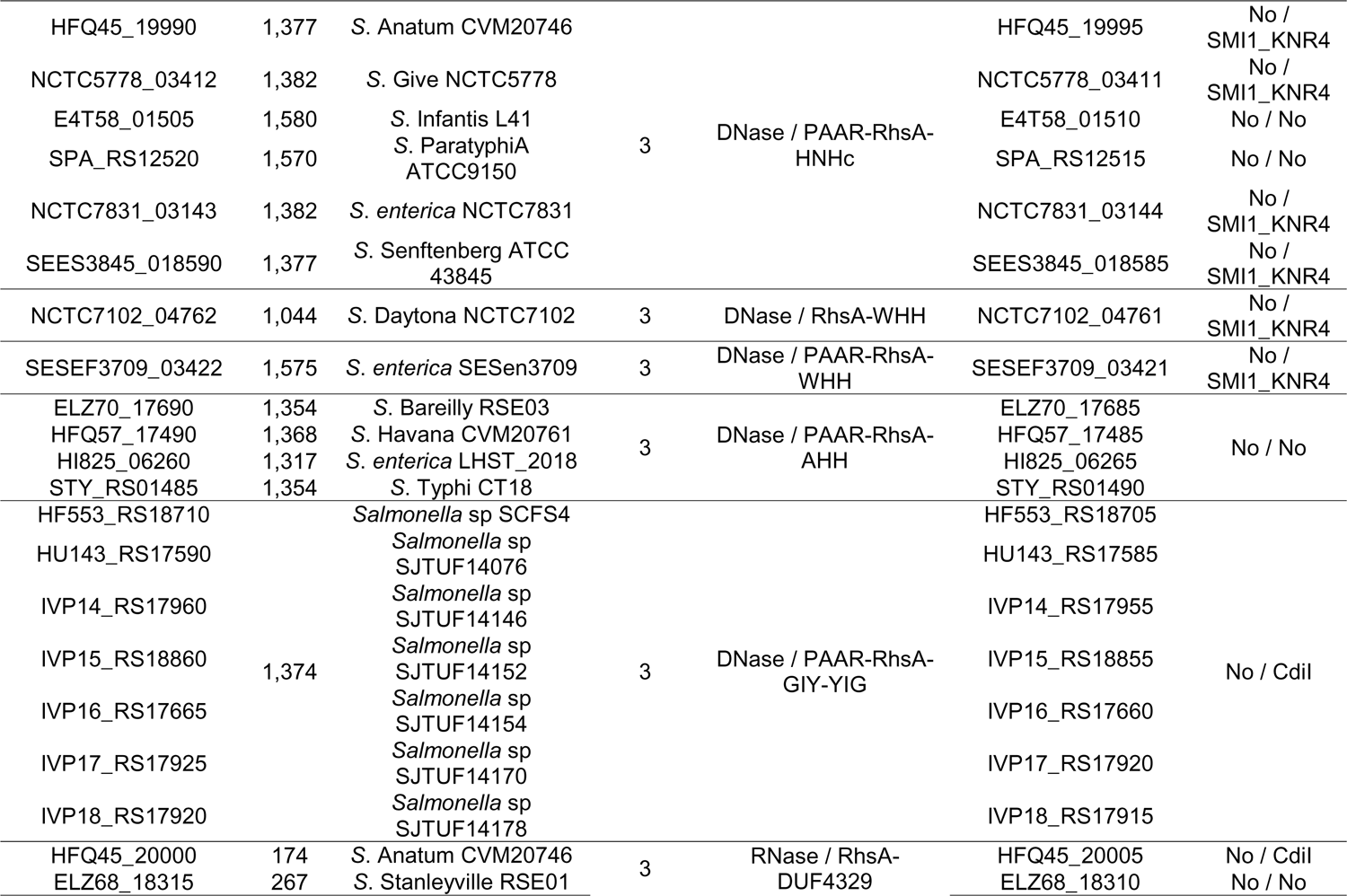
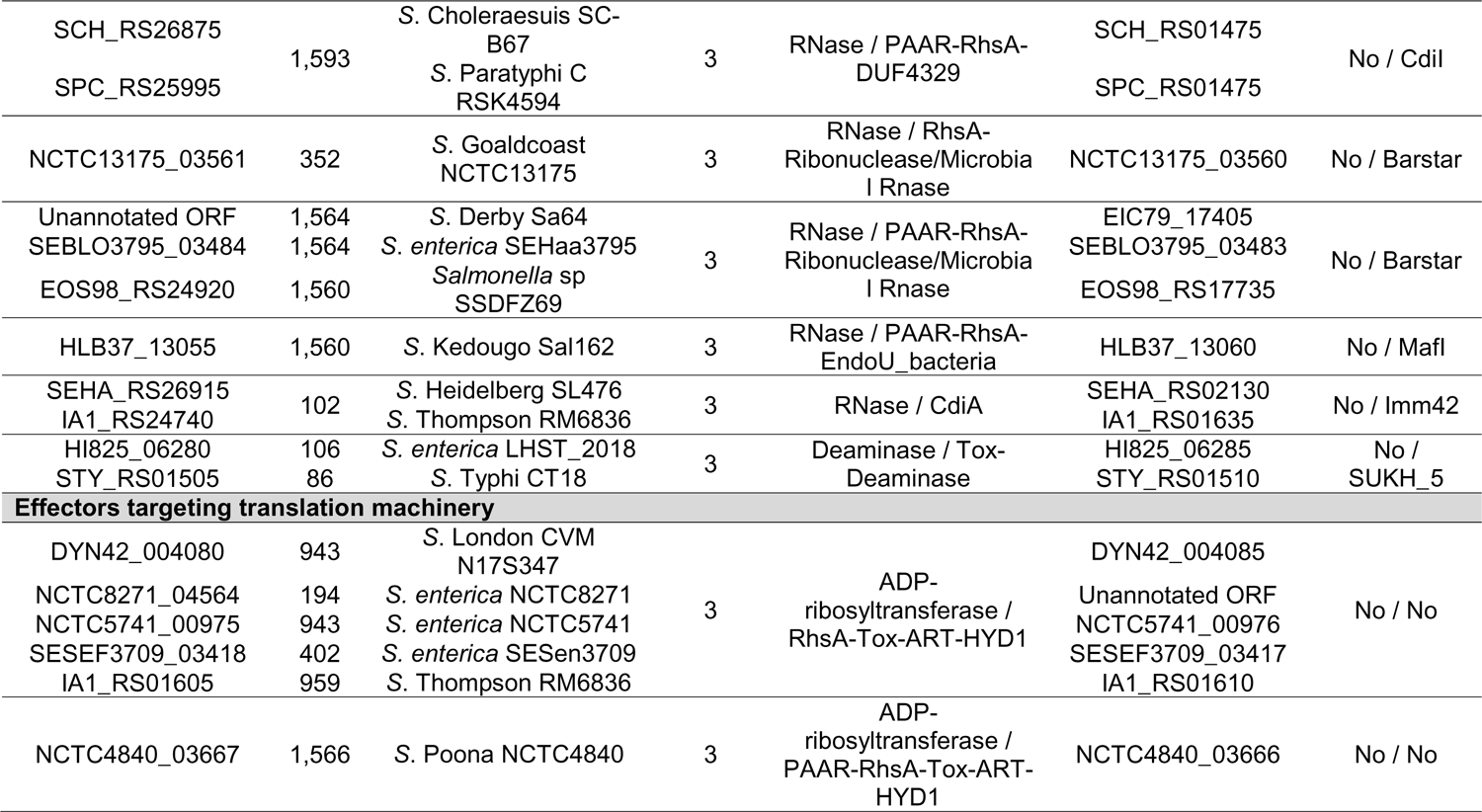
Novel putative T6SS effectors and cognate immunity proteins encoded in SPI-6 of *Salmonella enteric*.

### Putative T6SS cargo effectors with predicted peptidoglycan hydrolase activity are confined to VR1 and VR2

Our bioinformatic analysis identified 5 predicted T6SS cargo effectors with putative peptidoglycan hydrolase activity (**Table 1** and **Figure 4**). One effector corresponds to an unannotated ORF encoded within VR1. This ORF was identified in 30% (18/60) of the genomes analyzed and is localized between genes *tssH* and *tssB* (*ELZ70_17805* and *ELZ70_17795* ORFs in *S.* Bareilly strain RSE03) and is predicted to encode a 32 amino acids protein with a putative L-Ala-D-Glu-endopeptidase protein domain (**Figure 4**). This gene is predicted to be co-transcribed with a downstream unannotated ORF that encodes a 146 amino acids protein with a periplasmic-targeting signal peptide (**Table 1**), suggesting that this latter ORF encodes the cognate immunity protein of the novel candidate effector.

**Figure 4.**
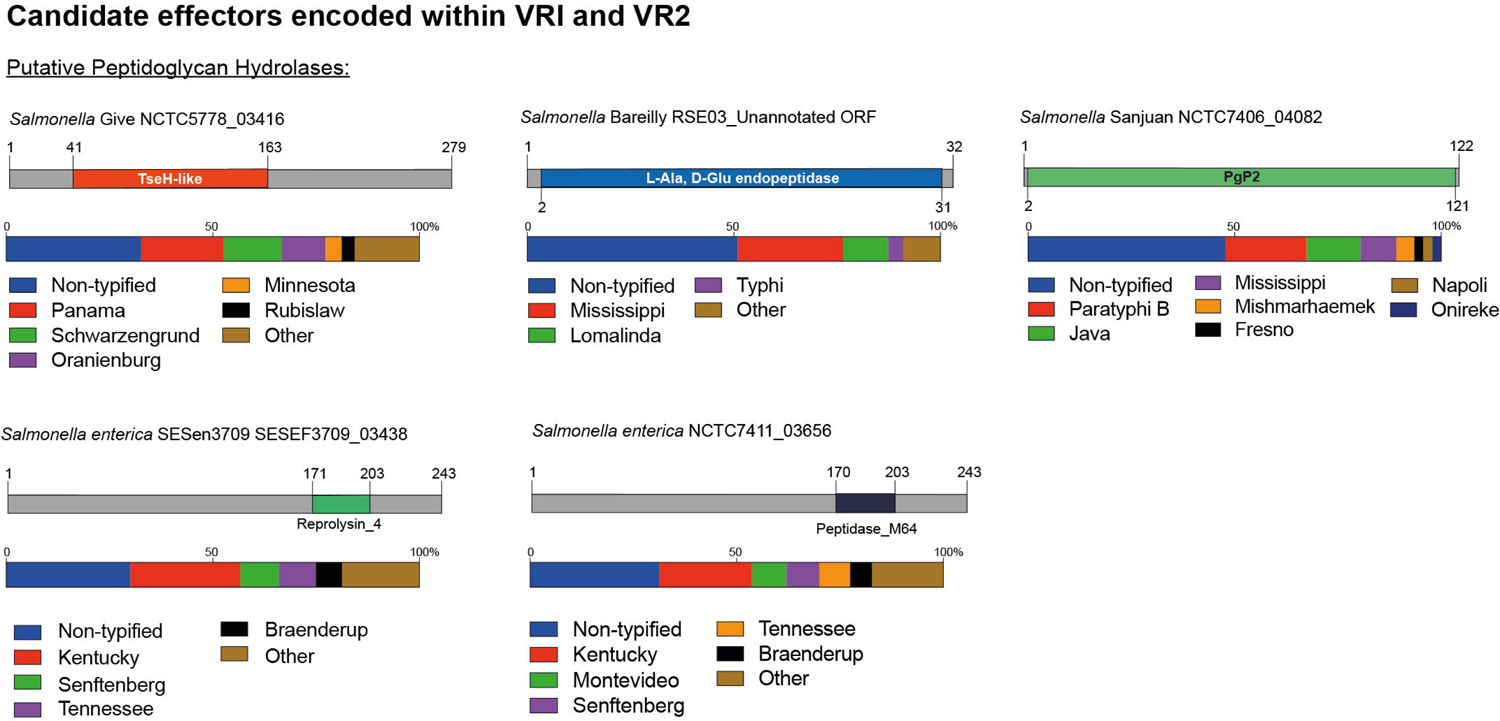
The variable regions 1 and 2 of the SPI-6 T6SS gene cluster encode 5 novel putative effectors. Schematic representation and distribution of novel putative effectors among *Salmonella* genomes. Predicted functional domains are show in different colors. Homologs for each candidate effector were identified by BLASTn analyses, as described in Materials and Methods.

In addition, our analysis identified four putative E/I modules encoded in VR2. The first putative effector (NCTC7406_04082 in *S*. Sanjuan strain NCTC7406) is a 122 amino acid protein that harbors a predicted PgP2 protein domain with putative L,D transpeptidase activity (**Table 1** and **Figure 4**). *NCTC7406_04082* is part of a bi-cistronic unit with *NCTC7406_04081*. This latter ORF encodes a 147 amino acid protein with a signal peptide targeting the periplasmic space that may correspond to its cognate immunity protein (**Table 1**). The second VR2 candidate effector (G9X22_18260 in *S*. Adjame strain 353868) is a 279 amino acids protein that harbors a putative amidase domain similar to the NlpC/P60 endopeptidase domain of the TseH T6SS effector of *Vibrio cholerae* (Altindis *et al*., 2015). This candidate effector is also encoded next to a putative immunity protein of 86 amino acids harboring a DUF4229 protein domain and 2 transmembrane helices that may target this protein to the periplasmic space (**Table 1**).

The third candidate effector (SESEF3709_03438 in *S*. *enterica* strain SESen3709) is a 243 amino acids protein that harbors a Reprolysin_4 domain with putative amidase activity (**Figure 4**). *SESEF3709_03438* is predicted to be part of a bi-cistronic unit with *SESEF3709_03437*, that encodes a putative cognate immunity protein with a signal peptide for periplasmic targeting.

The final candidate effector of VR2 corresponds to a 243 amino acid protein with a predicted M64 peptidase domain (NCTC7411_03656 in *S. enterica* strain NCTC7411) (**Table 1** and **Figure 4**). Our analysis also revealed that *NCTC7411_03656* is part of bi-cistronic unit with their respective putative immunity protein gene (*NCTC7411_03655* in *S. enterica* strain NCTC7411) (**Table 1**). In other serotypes, the putative immunity protein gene encodes a protein of 84-144 amino acids harboring a transmembrane domain that targets this protein to the periplasmic space (**Table 1**).

### Putative T6SS specialized effectors with polymorphic nuclease and ADP-ribosyltransferase toxin domains associated to Rhs proteins are restricted to the VR3

Our analysis revealed the presence of 18 candidate effectors encoded within the VR3 of SPI-6, including 16 of the Rhs family of proteins, 1 RNase and 1 deaminase. The size of the Rhs proteins ranged from 500 to 1,500 amino acids harboring different nuclease and ADP-ribosyltransferases domains (**Table 1** and **Figure 5**).

**Figure 5.**
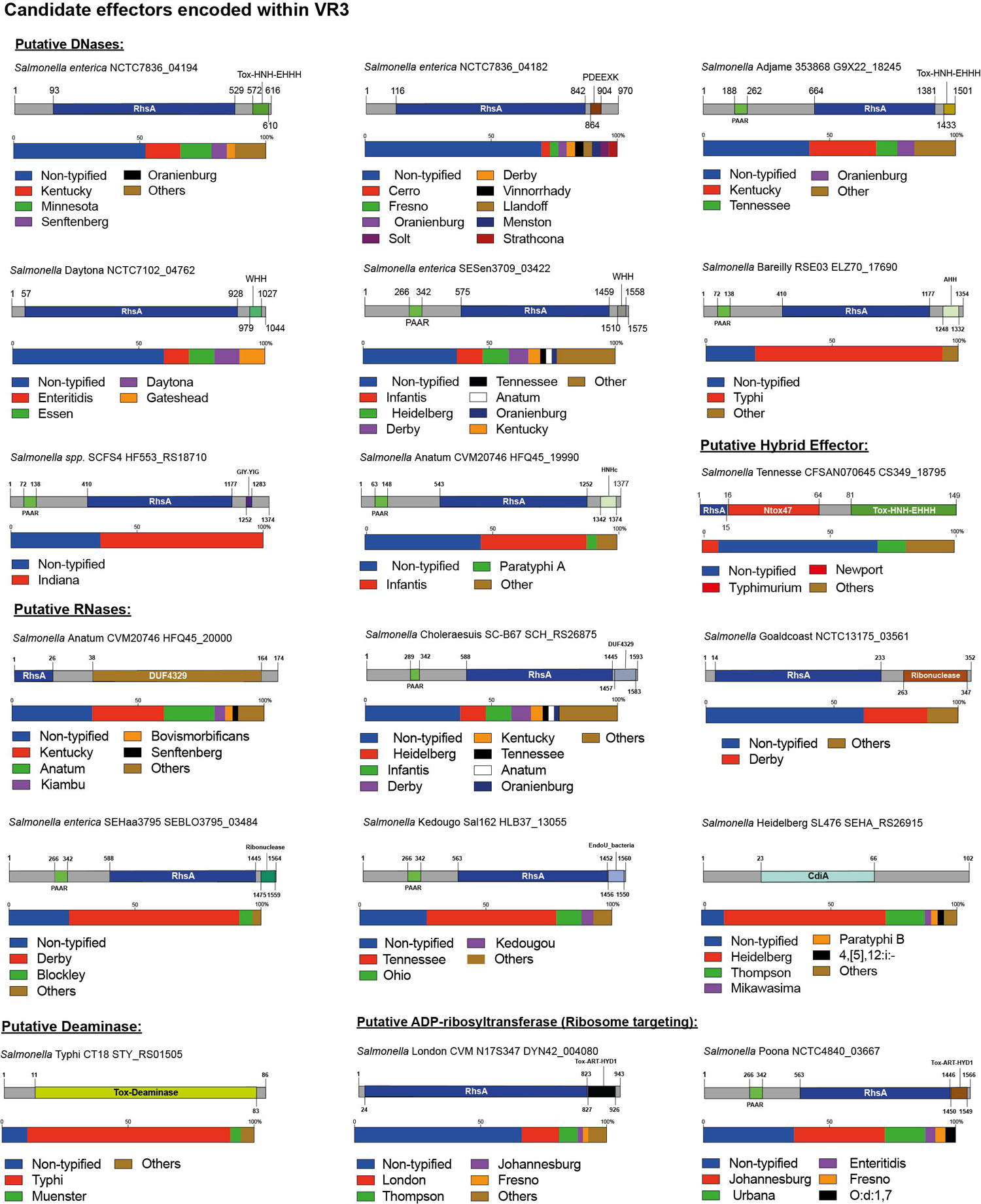
The variable region 3 of the SPI-6 T6SS gene cluster encodes 18 novel putative effectors. Schematic representation and distribution of novel putative effectors among *Salmonella* genomes. Predicted functional domains are shown in different colors. Homologs for each candidate effector were identified by BLASTn analyses, as described in Materials and Methods.

Eight of the 16 Rhs proteins harbored distinct C-terminal DNase nuclease domains, including domains of the HNH/ENDO VII superfamily of nucleases (IPR028048) such as WHH (IPR032869), Tox-HNH-EHH5H (IPR028048) or AHH (IPR032871), and nuclease domains of the GIY-YIG (IPR000305) and PDEEXK (IPR009362) families (**Figure 5**). In addition, 4 of these 8 candidates also harbored N-terminal PAAR motifs (IPR008727) (**Figure 5**). The presence of PAAR motifs suggests that these candidates correspond to specialized effector proteins. Each of these candidates were also predicted to be encoded in bi-cistronic units with ORFs encoding their respective immunity protein genes. Several of these proteins harbored domains previously found in cognate immunity proteins of bacterial toxin systems such as Imm50 (IPR028957), SMI1_KNR4 (PF09346) and CdI (IPR041256) (**Table 1**).

Our bioinformatics analyses also predicted 5 Rhs effectors with C-terminal extensions harboring different RNase protein domains (**Table 1** and **Figure 5**). These include Rhs proteins with Guanine-specific ribonuclease N1/T1/U2 (IPR000026), EndoU (IPR029501) and DUF4329 (IPR025479) domains. In addition, three of these proteins also harbored N-terminal PAAR motifs (IPR008727). The gene encoding each of these proteins was also predicted to be co-transcribed with genes encoding putative immunity proteins (**Table 1**). Remarkably, our analysis also identified a hybrid Rhs effector with predicted C-terminal RNase (Ntox47 domain) and DNase (Tox-HNH-EHHH) nuclease domains (CS349_18795 in *S*. Tennessee strain CFSAN070645). The gene encoding this protein is also predicted to be part of bi-cistronic unit with an ORF encoding a 129 amino acid protein with an Imm50 (IPR028957) domain. We also identified two putative Rhs effectors with a TOX-ART-HYD1 (pfam15633) ADP-ribosyltransferase domain, one of which also includes an N-terminal PAAR motif (NCTC4840_03667 in *S.* Poona strain NCTC4840). This protein shares 32% identity with STM0291, a recently described Rhs effector with an ART protein domain of *S.* Typhimurium named Tre^Tu^ (type VI ribosyltranferase effector targeting EF-Tu; Jurenas *et al*., 2022). The low percentage of sequence identity (**Figure S1**) suggests that this could be a divergent STM0291 homolog.

Finally, in VR3 we identified a putative effector with the CdiA RNase domain (IPR041620) not associated to Rhs elements (SEHA_RS26915 in *S*. Heidelberg SL476) (**Table 1**). In addition, we also identified a candidate effector harboring potential deaminase activity (STY_RS01505 in *S*. Typhi CT18). This effector is a small 36 amino acid protein with a TOX-deaminase domain of the BURPS668_1122 family (IPR032721) found in polymorphic toxin systems. The gene encoding this effector is predicted to be co-transcribed with an ORF encoding a putative immunity protein with a SMI1_KNR4 (PF09346) domain (**Table 1**).

### Genome-wide analysis of the distribution of SPI-6 T6SS effectors and candidate effectors in *Salmonella*

Identifying novel putative T6SS effectors encoded within VR1-3 of SPI-6 encourage us to determine the presence and distribution of the genes encoding these proteins across *Salmonella enterica*. The nucleotide sequence corresponding to each effector and candidate effector was used in BLASTn searches examining publicly available *Salmonella enterica* genome sequences deposited in the NCBI Database, and the distribution of each effector protein was determined (**Table S2**).

The analysis of the 9 T6SS effector proteins previously reported in the literature (i.e., Tae2, Tae4, Tge2, Tlde1, RhsA-HNHc, RhsA-Ntox47, PAAR-RhsA-Ntox47, Rhsmain-ART TreTu and Tox-URI2) and the 23 candidate effectors described in this study showed that they are widely and differentially distributed among *Salmonella* genomes (**Table S2**). Interestingly, we identified these effectors and candidates effector in many non-typified *Salmonella* strains (**Figure 6A**).

**Figure 6.**
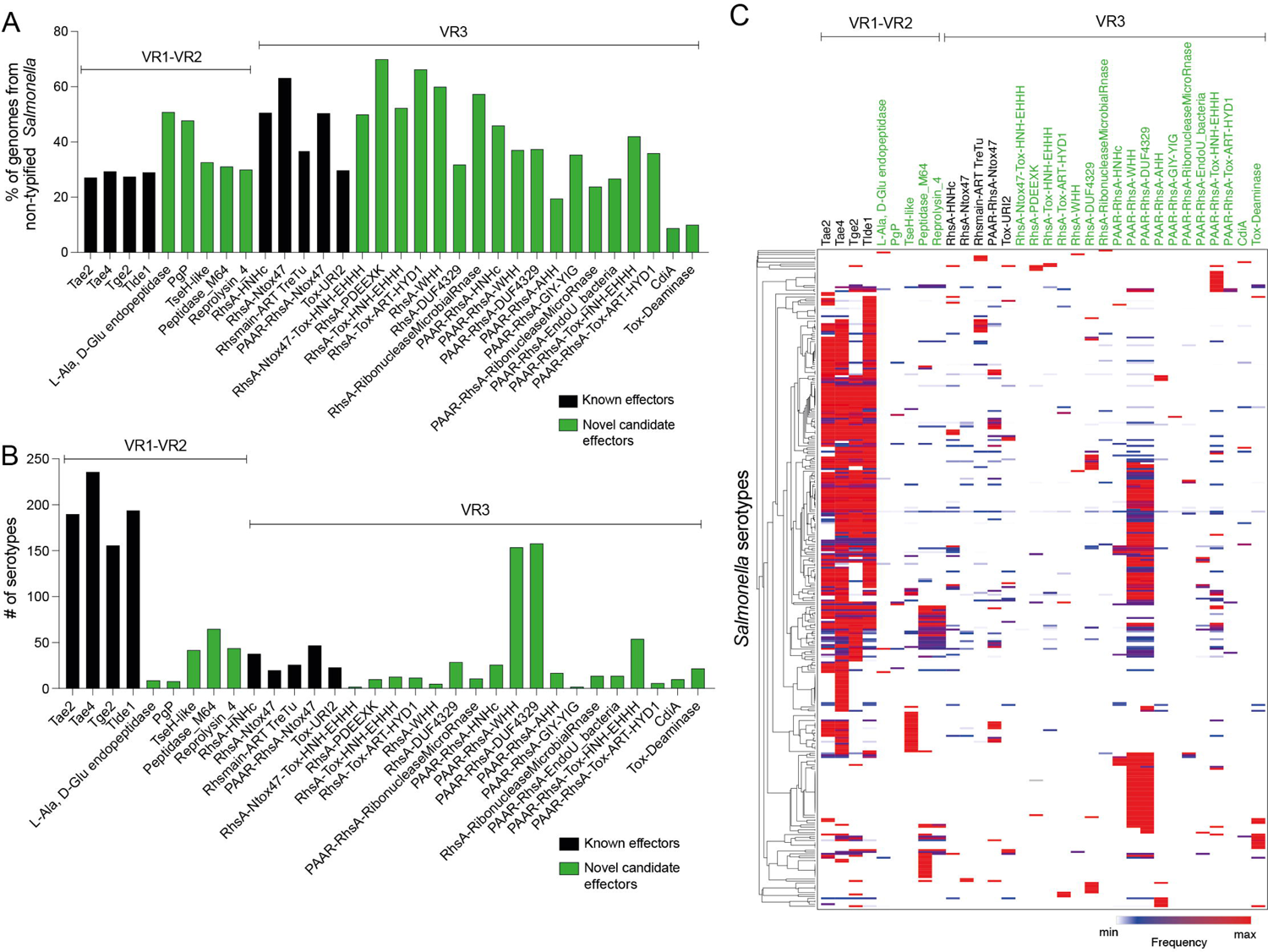
Prevalence of SPI-6 T6SS effectors and candidate effectors in *Salmonella*. Distribution of T6SS effectors and candidate effectors in non-typified (**A**) and serotyped (**B**) *Salmonella* strains. (**C**) Prevalence of T6SS effectors and candidate effectors encoded in the genome of 340 *Salmonella* serotypes. A hierarchical clustering analysis was performed using MORPHEUS, as described in Materials and Methods. Color code in the heatmap indicates the presence of a given effector (frequency) among all analyzed strains of a particular *Salmonella* serotype.

Some effector and candidate effectors were more widespread across different serotypes than others (**Figure 6B**). Within VR1 and VR2, the previously reported effectors Tae2, Tae4, Tge2, and Tlde1 were identified across 150-240 serotypes, while the five candidate effector proteins identified in this study were found in 5-50 distinct serotypes. A different scenario was observed for effectors and candidate effectors encoded within VR3. In this case, the previously reported effectors were identified in less than 50 serotypes, while some novel candidate effectors, such as PAAR-RhsA-WHH and PAAR-RhsA-DUF4329, were identified in over 150 serotypes. The distribution of each candidate effector in different *Salmonella* serotypes is highlighted in **Figure 4** and **Figure 5**.

Finally, we performed a hierarchical clustering analysis to gain further insight into the distribution of effector and candidate effectors identified in *Salmonella* genomes (**Table S3**). As shown in **Figure 6C**, the four *bona fide* effectors encoded within VR1-2 (Tae2, Tae4, Tge2 and Tlde1) were the most conserved across the genomes of different *Salmonella* serotypes. Nevertheless, these effectors are absent in the genome of several *Salmonella* serotypes, all of which include the genes encoding candidate effectors PAAR-RhsA-WHH and PAAR-RhsA-DUF4329 located within VR3. Furthermore, these candidate effectors are also widely distributed in the genome of several other *Salmonella* serotypes, suggesting that they play important roles in the biology of this pathogen.

## DISCUSSION

The T6SS has emerged as an important virulence and environmental fitness factor for *Salmonella* (Mulder *et al*., 2012, Pezoa *et al*., 2013, Pezoa *et al*., 2014, Sana *et al*., 2016, Sibinelli-Sousa *et al*., 2022, Xian *et al*., 2020, Blondel *et al*., 2010; Hespanhol *et al*., 2022). However, information regarding the complexity and diversity of effector proteins for each distinct *Salmonella* T6SS is still lacking. In this context, even though the T6SS_SPI-6_ has been shown to contribute to host colonization by *S*. Typhimurium and *S*. Dublin (Mulder *et al*., 2012; Pezoa *et al*., 2013; Pezoa *et al*., 2014; Sana *et al*., 2016) and to inter-bacterial competition of *S.* Typhimurium against the intestinal microbiota (Sibinelli-Sousa *et al*., 2022), only 9 effector proteins have been identified and characterized so far (Russell *et al*., 2012; Benz *et al*., 2013; Sana *et al*., 2016; Whitney *et al*., 2013; Sibinelli-Souza *et al*., 2020; Cobo *et al*., 2022; Koskiniemi *et al*., 2014; Amaya *et al*., 2022; Jurenas *et al*., 2022).

In this study, by means of bioinformatic and comparative genomic analyses, we identified a subset of 23 new SPI-6 T6SS candidate effectors, including peptidoglycan hydrolases, DNases, RNases, deaminases, and ADP-ribosyltransferases. Despite being well conserved, the SPI-6 T6SS gene cluster encodes a variable number of ORFs of unknown function restricted to three variable regions (VR1-3), that include the T6SS effectors previously identified in this species (Blondel *et al*., 2009). Notably, our analysis showed that every novel T6SS effector identified is encoded within one of these variable regions. An interesting observation was that all predicted peptidoglycan targeting effectors are confined to VR1 and VR2. The reason behind this observation remains unclear; however, it is possible that VR1 and VR2 are hot-spots for genes recombination during *Salmonella* evolution, but the lack of mobile genetic elements surrounding these regions does not support this hypothesis. Importantly, in addition to the 4 peptidoglycan targeting effectors reported so far (Tae2, Tae4, Tge2 and Tlde1) (Russell *et al*., 2012; Benz *et al*., 2013; Zhang *et al*., 2013; Sana *et al*., 2016; Whitney *et al*., 2013; Sibinelli-Sousa *et al*., 2020; Lorente-Cobo *et al*., 2022), we identified 5 candidate effectors encoded in VR1 and VR2 that degrade peptidoglycan, indicating that this macromolecule is a common target site for *Salmonella* T6SS effectors. Of note, the unannotated ORF encoded in VR1 is the first putative effector cleaving the link between L-Ala and D-Glu of the peptidoglycan peptide stems reported in *Salmonella* and shares homology to ChiX protein of *Serratia marcescens*, whose peptidoglycan hydrolase activity has been previously determined (Owen *et al*., 2018). This finding expands the peptidoglycan target sites exploited by *Salmonella* T6SS effectors against competing bacteria. On the other hand, the PgP2 and TseH-like candidate effectors have redundant peptidoglycan degrading functions with other *Salmonella* T6SS effectors. PgP2 is predicted to have the same L,D transpeptidase exchange activity reported for Tlde1 (Sibinelli-Sousa *et al*., 2020; Lorente-Cobo *et al*., 2022), replacing D-Ala by a non-canonical D-amino acid preventing the normal crosslink between mDAP and D-Ala. In addition, the TseH-like candidate effector is a NlpC/P60 endopeptidase (Altindis *et al*., 2015; Squeglia *et al*., 2019; Xu *et al*., 2010) predicted to cleave the covalent link between D-Glu and mDAP, as reported for Tae4 (Benz *et al*., 2013). These redundant functions suggests that the peptide stems are the main peptidoglycan target sites of *Salmonella* T6SS effectors, as the only identified effector targeting the glycoside bonds in this macromolecule corresponds to Tge2 (Whitney *et al*., 2013). The Reprolysin_4 domain found in some candidate effectors is present in zinc-binding metallo-peptidases harboring the binding motif HExxGHxxGxxH of family M12B peptidases. Of note, this motif is also present in the T6SS antibacterial effector SED_RS06335 with putative peptidoglycan hydrolase activity encoded in SPI-19 of *Salmonella* Dublin CT_02021853 (Amaya *et al*., 2022). The last candidate effector targeting the peptidoglycan identified in our study harbors the Peptidase_M64 protein domain that is also present in *Clostridium ramosum* IgA proteinase (Kosowska *et al*., 2002). The putative immunity proteins of Reprolysin_4 and Peptidase_M64 have a signal peptide and a transmembrane domain, respectively. This suggests that both candidate effectors target the bacterial periplasm.

On the other hand, the VR3 of the SPI-6 T6SS gene cluster encodes a wide variety of effector proteins including domains found in DNases, RNases, deaminases and ADP-ribosyltransferases. Interestingly, most of these domains are fused to the C-terminal of Rhs proteins contributing to diversify the molecular targets of T6SSs in *Salmonella*. This was not unexpected since we have previously shown that the VR3 of SPI-6 encodes a variable number of Rhs elements (Blondel *et al*., 2009; Amaya *et al*., 2022) and many Rhs proteins have C-terminal polymorphic endonuclease domains associated with T6SS effectors in *Salmonella* and other bacteria (Zhang *et al*., 2012, Koskiniemi *et al*., 2014; Amaya *et al*., 2022). It is known that Rhs proteins have YD-peptide repeats, which fold into a large β-cage structure that surrounds and protects the C-terminal toxin domain increasing T6SS secretion efficiency (Donato *et al*., 2020). This could explain why many T6SS effectors are associated to these elements.

Altogether, our work expands the repertoire of *Salmonella* T6SS effectors and provides evidence that the SPI-6 T6SS gene cluster harbors a great diversity of antibacterial effectors encoded in three variable regions. One interesting finding of our study is that peptidoglycan hydrolyzing effectors restricted to VR1 and VR2 are highly conserved in *Salmonella* genomes, while effectors targeting nucleic acids and the translation machinery encoded in VR3 are broadly distributed in *Salmonella* serotypes. This suggests that different repertoires of effectors could have an impact on the pathogenic potential and environmental fitness of these bacteria. Importantly, although this study increases the number of *Salmonella* antibacterial effectors against competing bacteria, we could not rule out that those encoded in VR3 may also target eukaryotic cells. This is an important knowledge gap, since no T6SS effector protein identified to date in *Salmonella* has been confirmed to target eukaryotic organisms, despite the clear contribution of *Salmonella* T6SSs to intracellular replication, survival and cytotoxicity inside the host immune cells (Mulder *et al*., 2012; Blondel *et al*., 2013; Schroll *et al*., 2019). Further research is required to address this issue. Finally, we are currently performing experimental work to confirm that each of the 23 candidates identified in our study correspond to *bona fide* T6SS effector proteins.

## AUTHOR CONTRIBUTIONS

CJB, FA, CAS and DP: conceptualization, formal analysis, validation, writing-original draft preparation, writing review and editing, resources, project administration, and funding acquisition. CJB and DP: methodology, investigation, and visualization. CAS and DP: supervision. All authors read and approved the final manuscript.

## SUPPLEMENTARY MATERIAL

**Table S1.** Dataset of *Salmonella* genomes retrieved from the Secret6 database.

**Table S2.** Distribution of T6SS effectors and candidate effectors in *Salmonella* genomes.

**Table S3.** Frequency of *Salmonella* strains of a particular serotype harboring each effector and candidate effector.

**Figure S1. Sequence alignment of effector proteins NCTC4840_03667 from S. Poona NCTC4840 and STM0291 from S. Typhimurium LT2.** A BLASTp alignment was performed using T-Coffee Expresso and visualized by ESPript 3. Identical amino acids are highlighted in boxes with a red background. Similar amino acids are highlighted in open boxes.

