## Supplementary figures and images for "Identification and distribution of novel candidate T6SS effectors encoded in *Salmonella* Pathogenicity island 6"

### Figure S1

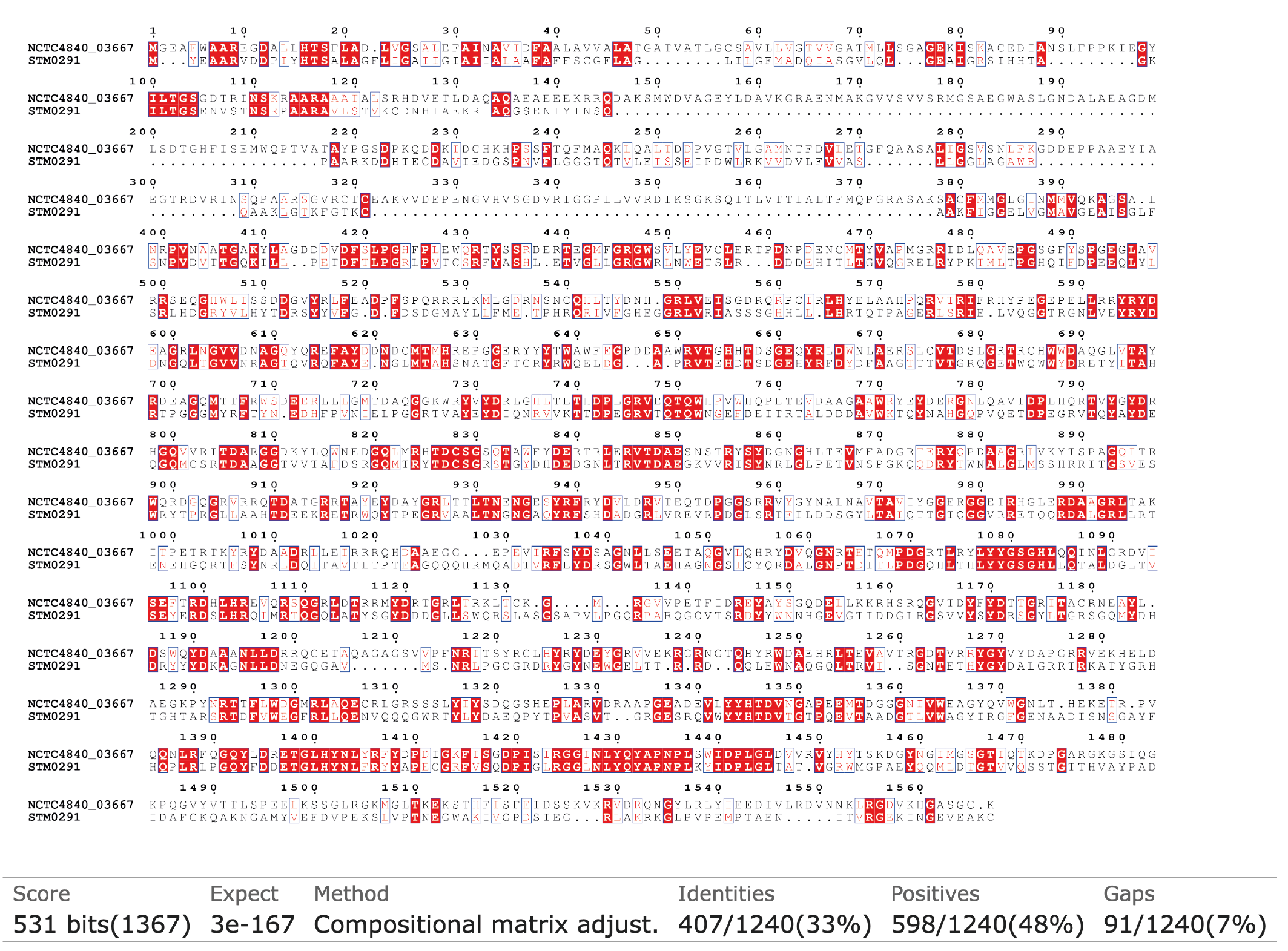
